# Determinants of early afterdepolarization properties in ventricular myocyte models

**DOI:** 10.1101/373266

**Authors:** Xiaodong Huang, Zhen Song, Zhilin Qu

## Abstract

Early afterdepolarizations (EADs) are spontaneous depolarizations during the repolarization phase of an action potential in cardiac myocytes. It is widely known that EADs are promoted by increasing inward currents and/or decreasing outward currents, a condition called reduced repolarization reserve. Recent studies based on bifurcation theories show that EADs are caused by a dual Hopf-homoclinic bifurcation, bringing in further mechanistic insights into the genesis and dynamics of EADs. In this study, we investigated the EAD properties, such as the EAD amplitude, the inter-EAD interval, and the latency of the first EAD, and their major determinants. We first made predictions based on the bifurcation theory and then validated them in physiologically more detailed action potential models. These properties were investigated by varying one parameter at a time or using parameter sets randomly drawn from assigned intervals. The theoretical and simulation results were compared with experimental data from the literature. Our major findings are that the EAD amplitude and takeoff potential exhibit a negative linear correlation; the inter-EAD interval is insensitive to the maximum ionic current conductance but mainly determined by the kinetics of I_Ca,L_ and the dual Hopf-homoclinic bifurcation; and both inter-EAD interval and latency vary largely from model to model. Most of the model results generally agree with experimental observations in isolated ventricular myocytes. However, a major discrepancy between modeling results and experimental observations is that the inter-EAD intervals observed in experiments are mainly between 200 and 500 ms, irrespective of species, while those of the mathematical models exhibit a much wider range with some models exhibiting inter-EAD intervals less than 100 ms. Our simulations show that the cause of this discrepancy is likely due to the difference in I_Ca,L_ recovery properties in different mathematical models, which needs to be addressed in future action potential model development.

**Author summary:** Early afterdepolarizations (EADs) are abnormal depolarizations during the plateau phase of action potential in cardiac myocytes, arising from a dual Hopf-homoclinic bifurcation. The same bifurcations are also responsible for certain types of bursting behaviors in other cell types, such as beta cells and neuronal cells. EADs are known to play important role in the genesis of lethal arrhythmias and have been widely studied in both experiments and computer models. However, a detailed comparison between the properties of EADs observed in experiments and those from mathematical models have not been carried out. In this study, we performed theoretical analyses and computer simulations of different ventricular action potential models as well as different species to investigate the properties of EADs and compared these properties to those observed in experiments. While the EAD properties in the action potential models capture many of the EAD properties seen in experiments, the inter-EAD intervals in the computer models differ a lot from model to model, and some of them show very large discrepancy with those observed in experiments. This discrepancy needs to be addressed in future cardiac action potential model development.

## INTRODUCTION

Under diseased conditions or influence of drugs, cardiac myocytes can exhibit early afterdepolarizations (EADs) [1,5]. EADs are depolarization events during the repolarizing phase of an action potential (AP), which are known to be arrhythmogenic [6-9]. Many experimental and computational studies have been carried out, which have greatly improved our understanding of the causes and mechanisms of EADs [10]. It is well known that EADs can occur in an AP when inward currents are increased and/or outward currents are reduced, a condition called reduced repolarization reserve [11]. Under this condition, L-type calcium (Ca^2+^) current (I_Ca,L_) can be reactivated to cause depolarizations in the repolarization phase of the AP to manifest as EADs. The importance of I_Ca,L_ reactivation for EAD genesis has been widely demonstrated in experiments [1,2] and computer simulations [12,13]. Recent studies [4, 14-16] using bifurcation theories have brought in additional mechanistic insights into the genesis of EADs, which show that EADs are oscillations originating via a supercritical or subcritical Hopf bifurcation and terminating via a homoclinic bifurcation, or via an unstable manifold of a saddle focus fixed point in the full AP dynamics [15]. Despite a great amount of studies on the genesis of EADs, less attention has been paid on the EAD properties, such as the EAD latency (the time from the upstroke of the AP to the upstroke of the first EAD), the inter-EAD interval, and the EAD amplitude. Although it is well-known that reactivation of I_Ca,L_ is required for EAD genesis, since EADs are a collective behavior arising from the interactions of many ionic currents, it is unclear how these ionic currents affect the EAD properties and what are the major determinants. For example, since I_Ca,L_ plays a critical role in the genesis of EADs, one would intuitively expect that increasing the maximum conduction of I_Ca,L_ might increase the amplitude of EADs, but as we show in this study that this is not case. On the other hand, understanding the EAD properties and their determinants is important for understanding the mechanisms of EAD-related arrhythmogenesis. For example, in an early experimental study [17], Damiano and Rosen showed that phase-2 EADs cannot propagate as premature ventricular complexes (PVCs) while phase-3 EADs can propagate as PVCs. This was also shown in our simulation studies [18-20]. Therefore, understanding what determine the EAD amplitude and takeoff potential may provide insights into EAD propagation to produce PVCs. If a PVC is a direct consequence of EAD propagation, then the EAD latency may provide information for the coupling interval between a sinus beat and the following PVC. EADs are also thought to be responsible for focal arrhythmias in the heart, and if this is true, then the oscillation frequency of EADs should be the same as the excitation frequency of ventricular arrhythmias. Furthermore, understanding the EAD properties and their determinants can also be important for the development of robust mathematical AP models. For example, we observed a discrepancy in inter-EAD interval between those from some widely used AP models and the experimental data. Experimental measurements in ventricular myocytes isolated from animal and human hearts almost exclusively show that the inter-EAD intervals are greater than 200 ms with few exceptions (Table I). However, many ventricular myocyte AP models show inter-EAD intervals much shorter than 200 ms [13,18,21-25], raising a question on what ionic current properties have been missed in these models.

In this study, we used bifurcation theories and computer simulations to systematically investigate the EAD properties and their major determinants. We first made theoretical predictions of the EAD properties using the 1991 Luo and Rudy (LR1) model [26] based on our previous bifurcation theory of EADs [4]. We then carried out computer simulations using physiologically more detailed ventricular AP models [18,20,24,27-29] to verify the theoretical predictions. In computer simulations, we also used parameter sets randomly drawn from assigned intervals so that large parameter ranges are explored to ensure generality of the simulation results. Theoretical and simulation results were compared with experimental results, and potential caveats of the current AP models were discussed.

## METHODS

### Action potential models

Computer simulations were carried out in single ventricular myocytes. The governing equation of the transmembrane voltage (V) for the single cell is

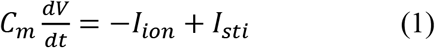

where I_ion_ is total ionic current density and I_sti_ the stimulus current density. C_m_ is the membrane capacitance which was set as C_m_=1 μF/cm^2^. We simulated six ventricular AP models: the 1991 Luo and Rudy (LR1) guinea pig model [26]; the 1994 Luo and Rudy (LRd) guinea pig model in a modified version [27]; the UCLA (H_UCLA_) rabbit model with modifications by Huang et al [20]; the 2004 ten Tusscher et al (TP04) human model [28]; the Grandi et al (GB) human model [24]; and the O’Hara et al (ORd) human model [29].

### Simulation methods

The differential equations were numerically solved using a first-order Euler method and the Rush and Larson method [30] for the gating variables with a fixed time step Δt=0.01 ms.

### Control parameters and parameter variations

For each model, a set of control parameters was used. The control parameter set is not the parameter set of the original model but a set we used for the AP to exhibit EADs. The major changes of parameters from the original models are either by increasing the maximum conductance of both I_Ca,L_ (was called slow inward current in the LR1 model, denoted as I_si_) and the slow component of the delayed rectifier potassium current (I_Ks_) or by increasing the maximum conductance of I_Ca,L_ but decreasing I_Ks_ or I_Kr_ (the rapid component of the delayed rectifier potassium current). The former corresponds to a normal myocyte (the original model) under isoproterenol while the later corresponds to the condition of long QT syndrome with isoproterenol. The specific changes of each model are detailed in the online Supplemental Information and the control APs exhibiting EADs are shown in Fig.S1.

To explore the effects of ionic current conductance on EAD properties in a wide parameter range, we varied the parameters in two ways. 1) We varied one parameter incrementally at a time but kept other parameters in their control values. The fold change of a specific parameter p is then defined as

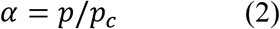

where p_c_ is the control value of p. 2) We randomly selected parameter sets, with each parameter drawn randomly from a uniform distribution in the interval (0.4p_c_, 1.6p_c_).

### Defining the EAD properties

We investigated three EAD properties in the AP models—amplitude, inter-EAD interval, and latency. As shown in Fig.1A, the EAD amplitude (A_EAD_) is defined as the difference between the takeoff voltage (V_takeoff_) and the peak voltage (V_peak_). The inter-EAD interval (T_EAD_) is defined as the time interval between the peaks of two consecutive EADs. The latency (L_EAD_) is defined as the time interval between the AP upstroke and the time when the 1st EAD takes off.

**Figure 1.**
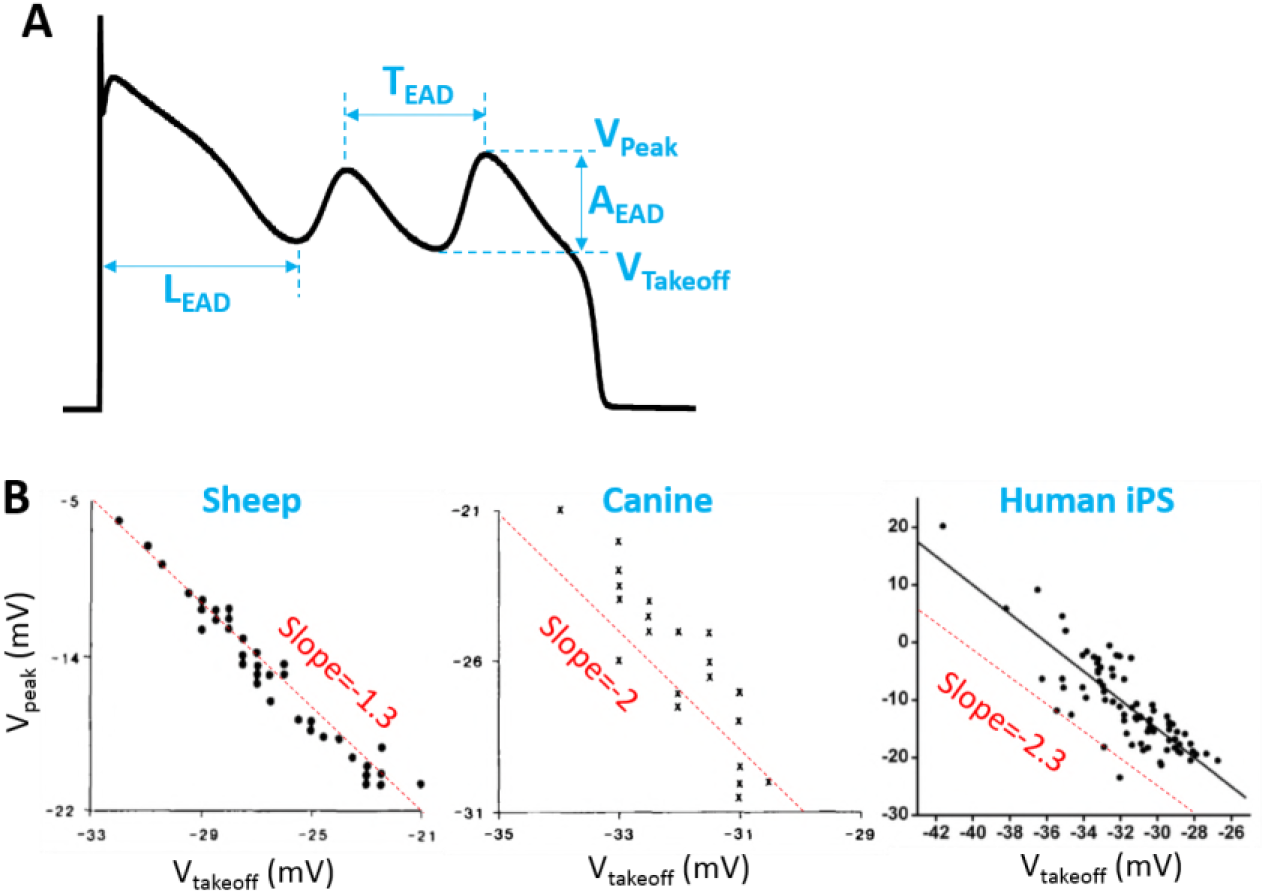
EAD properties. **A**. Definitions of A_EAD_, T_EAD_, L_EAD_, V_peak_, and V_takeoff_. **B**. V_peak_ versus V_takeoff_ from isolated sheep ventricular myocytes, canine ventricular myocytes [1,2], and human iPS cells [3]. The dashed straight lines were added as references for the slopes of the negative linear corrections of the data.

## RESULTS

### Experimentally observed EAD properties

Table I summarizes the EAD properties observed experimentally in isolated ventricular myocytes from literature survey [3, 31-51], which includes V_takeoff_, A_EAD_, T_EAD_, and L_EAD_. Based on this literature survey, we found that the V_takeoff_ is always above −50 mV except some of the mouse [51] and guinea pig [46,47] experiments. However, the EADs in the guinea pig experiments [46,47] were induced by injection of a constant inward current, which may behave differently from the ones occurring intrinsically. Therefore, the observed EADs in isolated ventricular myocytes are mainly phase-2 EADs. The maximum amplitude of the phase −2 EADs can be as large as 70 mV. It has also been observed in experiments that V_peak_ (thus A_EAD_) exhibits a negative linear correlation with V_takeoff_ (Fig.1B). As shown in Table I, T_EAD_ ranges from 200 ms to 500 ms except some of the mouse experiments [51]. L_EAD_ varies in a wide range, from as short as 30 ms to as long as 3 seconds.

**Table I.** Experimental EAD properties in isolated ventricular myocytes

| Reference (Figure) | Species | Lowest<br>$V_{\text{takeoff}}$ (mV) | Maximum<br>$A_{\text{EAD}}$ (mV) | $T_{\text{EAD}}$ (ms) | $L_{\text{EAD}}$ (ms) |
| --- | --- | --- | --- | --- | --- |
| [31] (Fig.2C) <sup>§</sup> | Human | -20 | 30 | ~ 250 | ~ 300 |
| [32] (Fig.10) | Human | -20 | 25 | 300 - 500 | 600 - 1000 |
| [33] (Figs.1A, 2A) | Human | -25 | 15 |  | 400 - 600 |
| [34] (Fig.2(b)) | Human | -15 | 25 | ~700 | ~1500 |
| [35] (Fig.6) <sup>&</sup> | Canine | -35 | 35 | ~ 270 | ~ 250 |
| [36] (Fig.1A) <sup>&</sup> | Canine | -30 | 25 | ~ 270 | ~ 270 |
| [36] (Fig.2A) <sup>&</sup> | Canine | -35 | 40 | 130 - 200 | ~ 160 |
| [37] (Figs.3A-C) | Canine | -25 | 25 | ~ 300 | ~ 300 |
| [38] (Fig.1B, 4A, 5A) | Canine | -25 | 30 | ~ 250 | ~ 300 |
| [39] (Fig.3) | Rabbit |  | 40 | ~ 300 | ~ 300 |
| [40] (Fig.6A) | Rabbit | -30 | 30 | ~ 300 | ~ 200 |
| [41] (Figs.1B, 2A) | Rabbit | -40 | 70 | ~ 400 | 400 - 500 |
| [42] (Figs.1, 3C-E) | Rabbit | -20 | 20 | 250 - 500 | 650 - 800 |
| [43] (Figs.1C, 6B) | Rabbit |  | 50 | 300 - 400 | 150 - 200 |
| [44] (Fig.1) | Rabbit | -10 | 10 | ~ 500 | ~ 450 |
| [45] (Figs.4B-C) | Guinea pig |  | 25 | ~ 200 | ~ 200 |
| [46] (Figs.2-4, 6, 8) <sup>#</sup> | Guinea pig | -40 | 70 | ~ 500 | 400 - 600 |
| [47] (Figs.2A, 3B) <sup>#</sup> | Guinea pig | -40 | 55 | ~ 200 | 200 - 1200 |
| [48] (Fig.5A) | Rat | -40 | 50 | 200 - 500 | 200 - 250 |
| [49] (Fig.6A) | Mouse | -45 | 50 | ~ 300 | ~ 3000 |
| [50] (Fig.6) | Mouse | -40 | 50 | ~170 | ~100 |
| [51] (Fig.4A), type I | Mouse | -40 | 45 | 50 - 90 | ~ 30 |
| [51] (Fig.4B), type II | Mouse | -55 | 40 | 40 - 90 | ~ 30 |

\*We estimated the following EAD properties: lowest takeoff potential, maximum EAD amplitude, inter-EAD interval, and EAD latency, using the AP traces shown in the figures as indicated in the parenthesis for each reference. We used the coordinates or the scale bars in the original plots to estimate these quantities. Therefore, the number shown in this table are not accurate measurements but just estimations.
<sup>§</sup>The time scale bar in Fig.2C in Ref. [31] is 300 ms instead of 150 ms per personal communication with Dr. G.-R. Li.
<sup>&</sup>The recordings in these studies [35,36] were not from truly isolated ventricular myocytes but from isolated small strips of M-cells.
<sup>#</sup>EADs were induced by constant inward current injection in these studies.

### Determinants of EAD amplitude and takeoff potentials

#### EAD amplitude in the LR1 model: Theoretical predictions and simulation results

We first investigated theoretically the relationship between A_EAD_ and V_takeoff_ based on bifurcation theories of EADs. In a previous theoretical study [4], we showed that phase-2 EADs are caused by a supercritical Hopf bifurcation and terminate at a homoclinic bifurcation in the fast subsystem of the AP, resulting in an oscillation with increasing amplitude and period (Figs.2 A and B). The oscillation amplitude

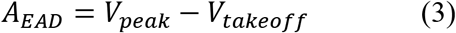

is bounded by a bell-shaped envelope centered at the upper quasi-equilibrium point (V_QES_, labeled as “Q” in Fig.2A). Based on the bell-shaped envelope of oscillation amplitude, one can approximate *V*_peak_ − *V*_QES_ ≈ (1 + *β*)(*V*_QES_ − *V*_takeoff_), then A_EAD_ can be written as:

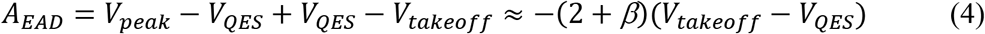

where β describes the degree of asymmetry of the bell-shaped envelope. For a symmetric oscillation (β=0), Eq. 4 predicts a linear relationship between A_EAD_ and V_takeoff_ with a slope −2. If one plots V_peak_ against V_takeoff_ as done in the plots of experimental data shown in Fig.1B, then Eq.4 is rewritten as

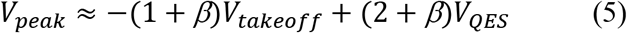

which predicts a slope −1 for an oscillation symmetric with respect to the equilibrium point. The experimental results shown in Fig.1B correspond to *β*=0.3, 1, and 1.3, respectively. A non-zero β indicates that the oscilaltions manifesting as EADs in the experiments are not symmetric to the quasi-equilibrium point and β vary from species to species. Note that Eq.5 is not obtained from a rigorous derivation but just an empirical observation based on the property of the dual Hopf-homoclinic bifurcation. The Hopf bifurcation shown in Fig.2A is a supercritical Hopf bifurcation, but previous studies [15,16] showed that a subcritical Hopf bifurcation could also be responsible for EAD genesis. In the latter case, Eqs.4 and 5 still hold based on the same reasoning.

**Figure 2.**
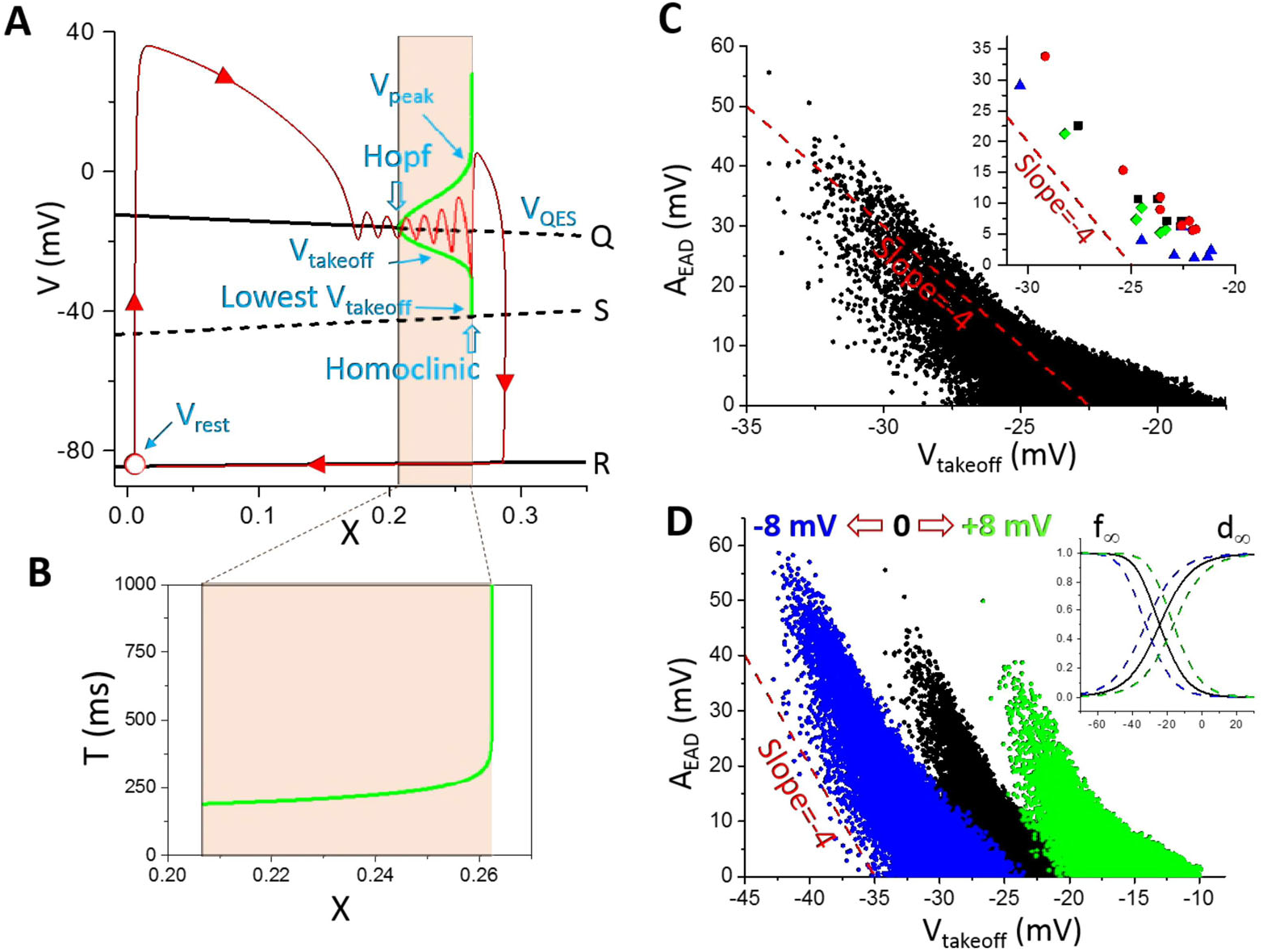
Bifurcation and EAD amplitude in the LR1 model. **A**. Bifurcations in the fast subsystem when treating the slow system (X) as a parameter. Q, S, and R are the three equilibria in the fast subsystem. The green bell-shaped envelope is the steady state oscillation amplitude (from V_takeoff_ to V_peak_) from the Hopf bifurcation point to the homoclinic bifurcation point. The lowest possible takeoff potential is at the homoclinic bifurcation point, which is around −41.5 mV. The red trace (arrows indicate the time course) is an AP from the whole system where X is a variable. Open red circle is the resting state. Note: the bifurcations and simulations shown in this panel and panel B were done without the presence of I_Na_. **B**. The steady-state oscillation period from the Hopf bifurcation point to the homoclinic bifurcation point. **C**. A_EAD_ versus V_takeoff_. The parameters were randomly drawn from the assigned intervals as described in Methods. For each parameter set, the amplitudes of all EADs in the AP were included. The inset shows representative A_EAD_ versus V_takeoff_ from individual APs distinguished by colors. **D**. Same as C but for shifted I_si_ kinetics (d_∞_ and f_∞_). The green data points are for d_∞_ and f_∞_ shifted 8 mV toward more negative voltages and the blue ones are for d_∞_ and f_∞_ shifted 8 mV toward more positive voltages. The inset shows the corresponding shifts. Panels A and B were replots from Tran et al [4]. The dashed straight lines in C and D are references for slopes.

We then carried out numerical simulations to investigate the relationship between A_EAD_ and V_takeoff_ in the LR1 model. For EADs from the same AP (the points with the same color in the inset of Fig.2C are from the same AP), their amplitudes and takeoff potentials approximately exhibit a linear relationship with slopes around −4 (the slope is around −3 if we plot V_peak_ against V_takeoff_ as in Fig.1B). We then plotted A_EAD_ against V_takeoff_ for all the EADs from the randomly drawn parameter sets. The data points are much more scattered, in particular when A_EAD_ is small. The lowest V_takeoff_ is around −34 mV with A_EAD_ ~ 55 mV. Based on the bifurcation analysis (Fig.2A), the lowest possible Vtakeoff in the LR1 model is around - 41.5 mV, and the numerical simulation results agree with the prediction of the bifurcation analysis. Since the EADs are caused by reactivation of I_Ca,L_ during the plateau phase, shifting the steady-state activation curve (d_∞_) and steady-state inactivation curve (f_∞_) simultaneously toward more negative or positive voltages results in almost the same shift of the A_EAD_ and V_takeoff_ relationship (Fig.2D). The maximum EAD amplitude becomes larger when the steady-state curves are shifted toward more negative voltages.

To investigate how A_EAD_ is affected by the maximum conductance and kinetics of different ionic currents, we varied one parameter at a time while maintaining other parameters at their control values. Fig.3A shows A_EAD_ versus the fold of the control G_si_ (*α* as defined in Eq.2) for EADs in the AP (note: I_Ca,L_ is denoted as I_si_ in the LR1 model and G_si_ is the maximum conductance). When *α* is smaller than 0.658, no EADs occur. When *α* is between 0.658 and 0.74, one EAD appears in the AP. As G_si_ increases, more EADs appear in the AP. When *α* is greater than 1.2, there are 10 EADs in the AP. Since I_si_ is an inward current, one may also anticipate that increasing I_si_ would progressively increase the EAD amplitude and APD. However, this is not the case. For example, increasing *α* from 0.66 to 0.74, the EAD amplitude decreases quickly from around 35 mV to around 10 mV and the APD also becomes shorter (see the APs in Fig.3B). As *α* is slightly above 0.74, a second EAD with a large amplitude (>30 mV) suddenly appears in the AP and also abruptly increases the APD (see the APs in Fig.3C). Note that the parameters for the blue and magenta traces differ slightly (a 0.3% difference), and the two traces are almost identical until at the very end of phase-2 where one repolarizes (exits the basin of attraction of the limit cycle) and the other depolarizes (retains in the basin of the limit cycle) to result in an EAD, a typical all-or-none behavior. The amplitude of this EAD decreases quickly and the APD also decreases as G_si_ increases (see the APs in Fig.3D) until another new EAD appears in the AP. This process repeats as G_si_ increases. Therefore, as shown in Fig.3A, the number of EADs in an AP increases progressively with G_si_, but the amplitude of a specific EAD decreases with G_si_ until a new EAD (all-or-none) suddenly appears in the AP with a maximum amplitude. In other words, the maximum EAD amplitude always occurs when a new EAD appears in the AP. Before a new EAD occurs, the amplitudes of all EADs in an AP are relatively small (such as the EADs in the black AP in Fig.3B). The overall maximum EAD amplitude does not exhibit an apparent change against G_si_ (remained at around 35 mV for *α* changed from 0.65 to 1.2).

**Figure 3.**
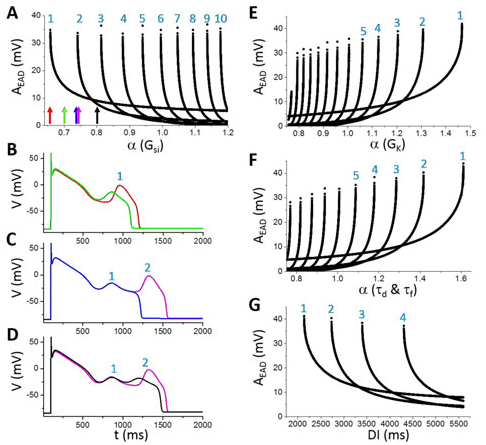
Dependence of EAD amplitude on ionic currents in the LR1 model. **A**. A_EAD_ versus fold of control G_si_ [labeled as α(G_si_)]. The colored arrows mark the α(G_si_) values for the traces shown in B-D: red, α=0.658; green, α=0.7; blue, α=0.74; magenta, α=0.7425; and black, α=0.8. The numbers mark the EAD order in an AP as indicated in B-D. For example, in B, there is only one EAD in both APs, then the EAD is labeled as the 1^st^ EAD. In C, a new EAD appears in the magenta AP, and this EAD is labelled the 2^nd^ EAD while the old one is labeled as the 1^st^ EAD. The umber increases as more EADs appear in the AP. **B**. APs with one EAD as α indicated by the red and green arrows in A. **C**. APs in the transition from one EAD to two EADs as α indicated by the blue and magenta arrows in A. **D**. APs with two EADs as α indicated by the magenta and black arrows in A. **E**. A_EAD_ versus fold of control G_K_. **F**. A_EAD_ versus fold of control τ_d_ and τ_f_. In these simulations, τ_d_ and τ_f_ are multiplied by the same α. **G**. A_EAD_ versus DI. α(G_si_)=0.95 and other parameters are their control values.

Increasing the conductance of the time-dependent outward K^+^ current (I_K_) decreases the number of EADs in the AP, but increases the EAD amplitude (Fig.3E), which is the opposite to the effects of increasing I_si_. As G_K_ increases, the amplitude of the last EAD reaches maximum immediately before its disappearance from the AP. Slowing the activation and inactivation time constants reduces the number of EADs in an AP but increases the maximum EAD amplitude (Fig.3F). The dependence of EAD amplitude on diastolic interval (DI) is similar to changing a conductance (Fig.3G), i.e., decreasing DI is equivalent to increasing G_K_.

To more closely link the dual Hopf-homoclinic bifurcation to the behaviors of EAD amplitude shown in Fig.3, we compared the bifurcations of the fast subsystem and the EADs of the whole system in the same way as in Fig.2A. Since changing the maximum conductance of ionic currents not only changes the EAD behavior but also changes the bifurcation of the fast subsystem, to only change the EAD properties but not the bifurcations of the fast subsystem, we changed the time constant of X-gate variable, τ_X_. This is because the bifurcation diagram is obtained by treating X as a parameter and thus the bifurcation will remain the same for any τ_X_. Fig.4A plots A_EAD_ versus the fold change of τ_X_, showing that increasing τ_X_ increases the number of EADs in the AP and gives rise to the same A_EAD_ behavior as changing the maximum conductance shown in Fig.3. Fig.4B shows three APs with α indicated by the colored arrows in Fig.4A. At the red arrow (the red AP trace in Fig.4B), there are four EADs in the AP. Increasing τ_X_ slightly (the blue arrow in Fig.4A and the blue AP trace in Fig.4B), a new EAD (the 5^th^ EAD) with a much larger amplitude appears in the AP. As shown in Fig.4C, the new EAD in the blue race takes off right before the homoclinic bifurcation while the red trace just passes the homoclinic bifurcation point during its 4^th^ EAD. The reason is that X grows slightly slower due to a slightly larger τ_X_ for the blue trace so that the 4^th^ EAD ends just right before the homoclinic bifurcation, allowing a new EAD to take off. Increasing τ_X_ further quickly reduces the EAD amplitude of the 5^th^ EAD because its takeoff point moves away from the homoclinic bifurcation point (open arrows in Fig.4D) due to a slower X growth caused by a larger τ_X_.

**Figure 4.**
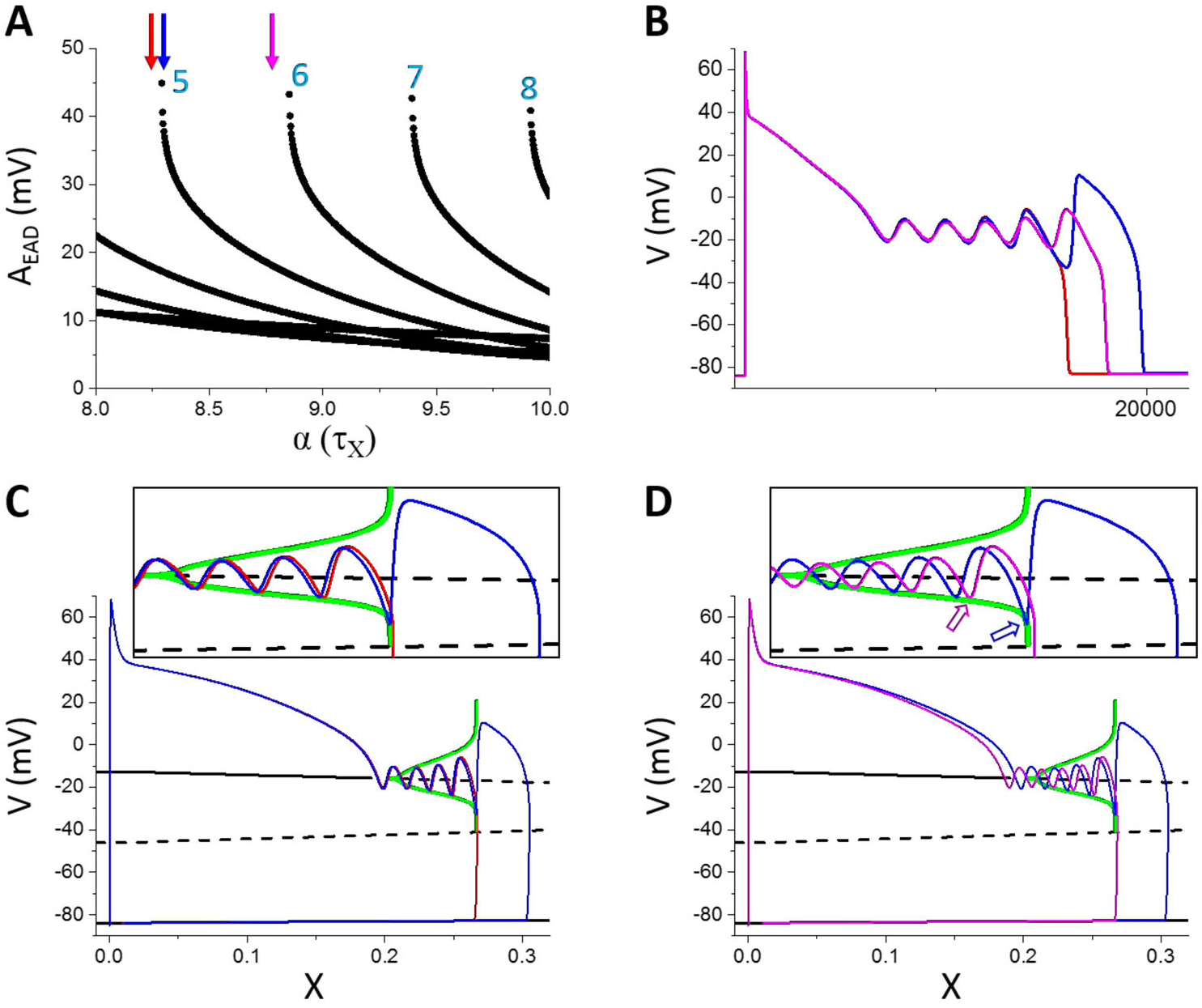
Linking the dual Hopf-homoclinic bifurcation to the EAD amplitude behavior. **A**. A_EAD_ versus fold change (α) of τ_X_. **B**. APs correspond to the parameters indicated by the arrows with the same colors. Red: α=8.24; Blue: α=8.29; Magenta: α=8.78. **C**. Bifurcation in the fast subsystem, the same as in Fig.2A except that I_Na_ was present. The blue and red traces are V versus X for the corresponding blue and red APs in B. Inset is the blowup of the EAD window. **D**. Same as C but for the blue and magenta APs in B. Open arrows in the inset mark the takeoff location of the 5^th^ EAD with respect to the homoclinic bifurcation point. Note that both APs exhibit 5 EADs but the ones in the magenta AP take off at smaller X values.

#### EAD amplitude in physiologically more detailed models

To further assess the theoretical predictions and simulation results from the LR1 model, we carried out simulations using physiologically more detailed AP models. Fig.5 shows A_EAD_ versus V_takeoff_ for all six models we simulated. The negative linear correlation holds roughly for all AP models while the slopes varied from −2 to −5 (corresponding to slopes ranging from −1 to −4 in plots of V_peak_ versus V_takeoff_). Shifting the steady-state activation and inactivation curves of I_Ca,L_ results in roughly the same shift in the A_EAD_ and V_takeoff_ relationship, indicating that the I_Ca,L_ reactivation is responsible for EADs in all the models. The maximum EAD amplitude and the lowest detectable takeoff potential also vary largely from model to model. The maximum EAD amplitude (without the shifts in I_Ca,L_ kinetics) of the TP04 model is the smallest (<35 mV) while that of the ORd model is the largest (~ 90 mV) among the six models. The lowest V_takeoff_ of the TP04 model is the highest (~ −20 mV) while that of the H_UCLA_ model is the lowest (~-45 mV) among the six models. Although the EAD amplitude properties vary largely from model to model, they generally agree with the experimental data shown in Fig.1B and Table I.

**Figure 5.**
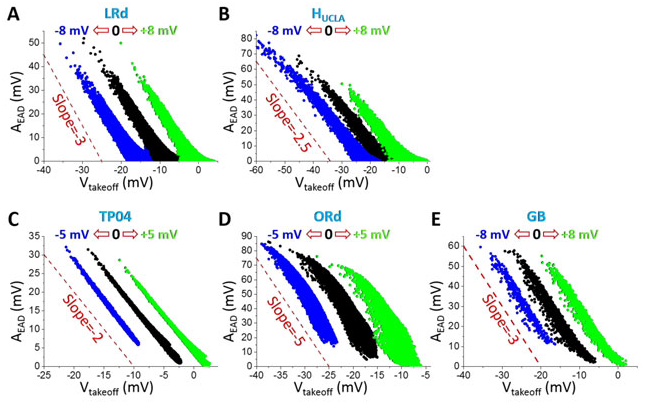
A_EAD_ versus V_takeoff_ in the physiologically more detailed models. **A**. LRd. **B**. H_UCLA_. **C**. TP04. **D**. ORd. **E**. GB. Parameters were randomly drawn from assigned intervals (as described in Methods). A_EAD_ was measured for all EADs in an AP. Black dots are A_EAD_ for control steady-state activation and inactivation curves of I_Ca,L_; blue dots are for both curves being shifted toward negative voltage; and green dots are for both curves being shifted toward more positive voltages. The voltage shifts are indicated by the open arrows in each panel in the same way as in Fig.2D. Dashed lines are reference lines for the slopes.

We investigated the effects of the maximum conductance of the major ionic currents on A_EAD_ for all six models, including I_Ca,L_ (Fig.S2), I_Ks_ (Fig.S3), I_Kr_ (Fig.S4), I_K1_ (Fig.S5), I_NCX_ (Fig.S6), I_NaK_ (Fig.S7), I_NaL_ (Fig.S8), and I_to_ (Fig.S9). The effects of the inactivation time constant of I_Ca,L_ were also shown (Fig.S10). The general observations are the same as those from the LR1 model, i.e., increasing an inward current increases the number of EADs in the AP, decreases A_EAD_ until a new EAD suddenly appears in the AP at which A_EAD_ becomes maximum. Increasing an outward current decreases the number of EADs in the AP, and increases A_EAD_ and A_EAD_ of the last EAD in the AP reaches maximum before it suddenly disappears from the AP. However, there are some exceptions. For example, increasing the maximum conductance of I_to_ can either promote or suppress EADs (Fig.S9). In both the LRd model and the ORd model, increasing I_to_ promotes EADs (more number of EADs in the AP), but suppresses EADs in the TP04 model and GB model. For the H_UCLA_ model, increasing I_to,s_ promotes EADs, while increasing I_to,f_ first promotes EADs but then suppresses EADs. Since I_to_ is an outward current, it is generally known that it suppresses EADs [52]. However, recent studies have demonstrated that I_to_ can also promote EADs [43,53]. Whether I_to_ promotes or suppresses EADs depends on its magnitude, speed of inactivation, and its pedestal component. Slow inactivation or large pedestal current tends to suppress EADs [53]. Another exception is I_NCX_. Increasing I_NCX_ promotes EADs in all the models except in the TP04 model in which EADs may also be suppressed by increasing I_NCX_ (see Fig.S6).

### Determinants of inter-EAD interval

As shown in Table I, T_EAD_ in isolated ventricular myocytes is typically from 200 ms to 500 ms. In this section, we investigate T_EAD_ and its determinants in the AP models. We first used the LR1 model for theoretical treatments and then used the physiologically more detailed models to verify the theory.

#### Inter-EAD interval in the LR1 model: Theoretical predictions and simulation results

Based on Tran et al [4], EADs arises from a Hopf bifurcation in the fast subsystem of the LR1 model. Here we first calculate the period of oscillation at the Hopf bifurcation point analytically and then compare the period of oscillation to the inter-EAD interval. The eigenvalue λ of the Jacobian (Eq.2 in Tran et al [4]) satisfies the following equation:

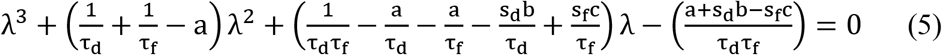

where τ_d_ and τ_f_ are the time constant of the d-gate and the f-gate of I_si_, respectively; s_d_ and s_f_ are the slopes of the steady-state activation and inactivation curves, respectively; and *a*, *b*, and *c* are derivatives of certain functions (see Tran et al [4] for definitions of these quantities). These quantities are their values at the steady state. At the Hopf bifurcation point, λ_1,2_ = ±iω, satisfying (λ − iω)(λ + iω)(λ − λ_3_) = 0, i.e.,

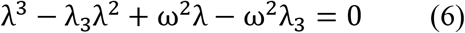

Comparing the coefficients of the Eqs. (5) and (6), one oobtains:

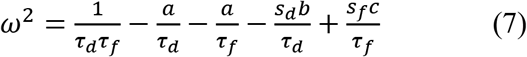

To calculate ω, one needs to define the Hopf bifurcation point. In the LR1 model, the slow subsystem is the *X*-gate which is treated as a parameter for the stability analysis of the fast subsystem. For a given *X* value, one can easily obtain the steady-state voltage V_QES_ and thus the steady-state values of all other gating variables. From Eqs. (5) and (6), one also has: 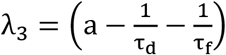 and 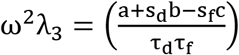 This leads to:

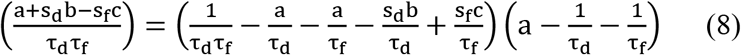

Defining

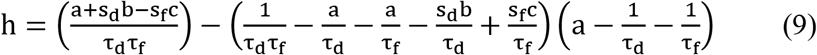

then at the Hopf bifurcation point, h = 0. Moreover, at the vicinity of the Hopf bifurcation point, λ_1,2_ = ε ± iω, satisfying (λ − ε − iω)(λ − ε + iω)(λ − λ_3_) = 0. Following the same analysis above, one obtains:

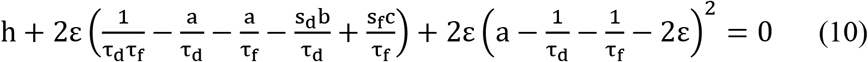

Since ε<0 before the Hopf bifurcation and ε>0 after the Hopf bifurcation, thus h>0 before the Hopf bifurcation and h<0 after the bifurcation. Therefore, by calculating h using Eq.9 and scanning *X* from 0 to 1, one can determine the Hopf bifurcation point. After the Hopf bifurcation point is determined, one can then calculate the oscillation period of the limit cycle of the fast subsystem at the Hopf bifurcation using Eq.7. Fig.6 shows the oscillation period versus different parameters using the theoretical approach (open red symbols). The oscillation period decreases slowly with increasing G_si_ (Fig.6A), has almost no change with G_K_ (Fig.6C) and G_K1_ (Fig.6D), but increases quickly with slowing the inactivation time constant of I_si_ (a 2-fold change in τf resulted in roughly a 2-fold change in the oscillation period, Fig.6E).

**Figure 6.**
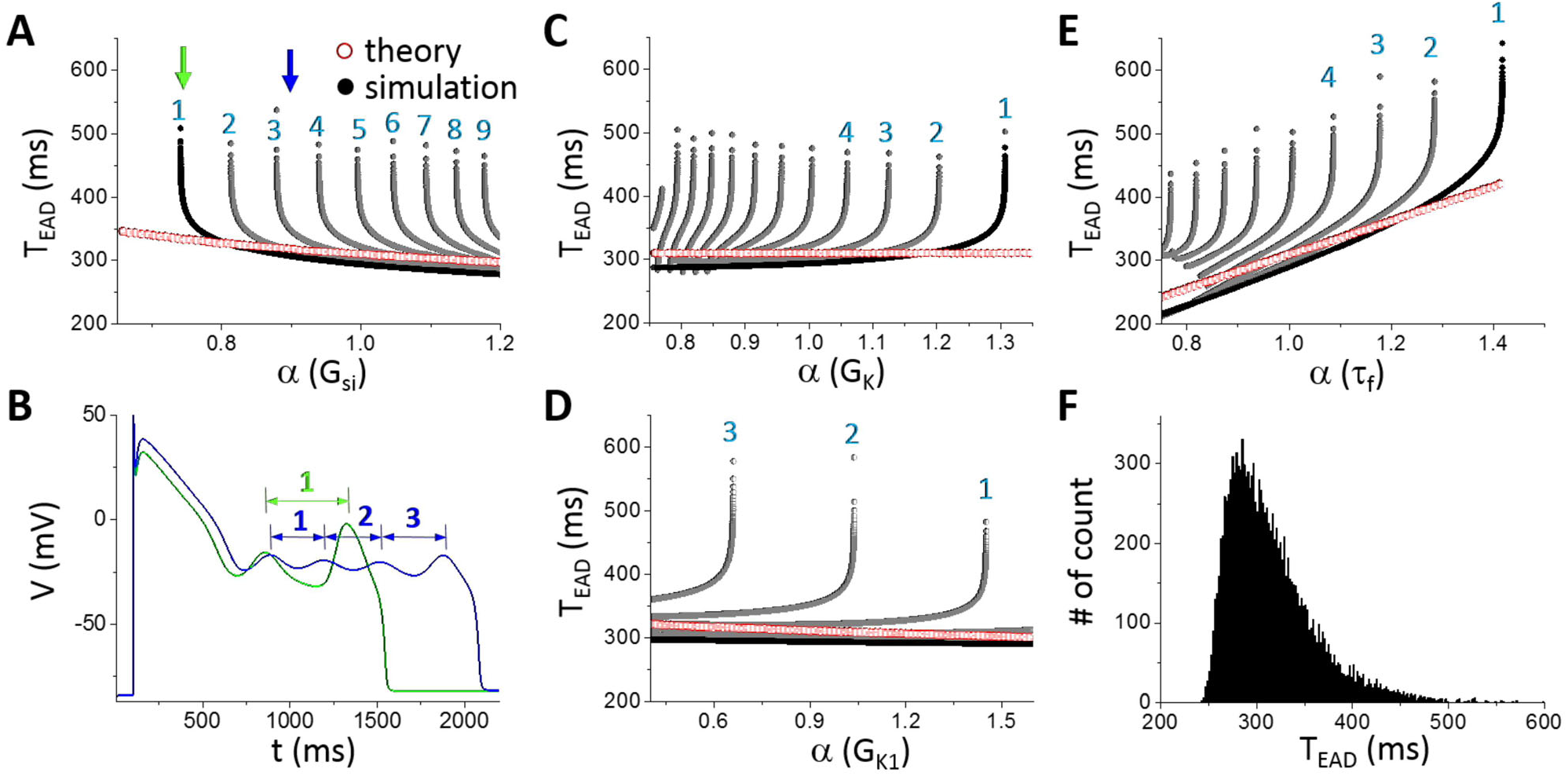
Theoretical predictions and simulation results of T_EAD_ using the LR1 model. **A**. T_EAD_ versus G_si_. The numbers mark the order of T_EAD_ as defined in B. **E**. AP traces for G_si_ indicated by the two arrows in A with two EADs and one T_EAD_ in the green AP (α=0.741) while four EADs and three T_EAD_ in the blue AP (α=0.9). **C**. T_EAD_ versus GK. **D**. T_EAD_ versus GK1. **E**. TEAD versus τ_f_. **F**. Histogram of T_EAD_ obtained from a large number of simulations using random selected parameter sets and all T_EAD_ in an AP. Control τ_f_ (α=1) was used. The total number of T_EAD_ is 13317 in the histogram.

For comparison, we carried out simulations of the LR1 model using the same parameters. We plotted all T_EAD_ in an AP as we did for A_EAD_ with the ordering of T_EAD_ in an AP shown in Fig.6B. Similar to A_EAD_, when a new EAD suddenly appears in or disappears from the AP, the appearing or disappearing T_EAD_ is the longest. As shown in Fig.6A, as G_si_ increases, an EAD suddenly appears in the AP (green arrow in Fig.6A and green AP in Fig.6B) which results in a long T_EAD_. As G_si_ increases further, this T_EAD_ decreases quickly toward the oscillation period of the limit cycle in the fast subsystem predicted from the theory, and becomes insensitive to G_si_. This T_EAD_ behavior repeats once a new EAD occurs in the AP as G_si_ increases. The fast decaying phase is a result of the homoclinic bifurcation, in which the oscillation period of the limit cycle decreases quickly as the system is away from the homoclinic bifurcation point (see Fig.2B). In other words, when a new EAD first appears in an AP, it is always the one closest to the homoclinic bifurcation, and thus the A_EAD_ is the largest and T_EAD_ the longest (see Fig.4). As the takeoff of this EAD is further away from the homoclinic point, and the corresponding T_EAD_ (e.g., from the blue AP to the magenta AP in Fig.4B) quickly approaches to those of the EADs prior to it, roughly the oscillation period of the limit cycle of the fast subsystem at the Hopf bifurcation. Therefore, changing G_si_ has only a small effect on T_EAD_ except when an EAD appears in or disappears from the AP. Changing the maximum conductance of an outward current exhibits the same behavior (Figs.6 C and D). However, T_EAD_ increases with the time constant τ_f_ of the I_Ca,L_ inactivation gate more sensitively than with the maximum conductance of the ionic currents (Fig.6E), which also agrees with the theoretical prediction. This indicates that besides the dual Hopf-homoclinic bifurcation, I_Ca,L_ inactivation and recovery (although τ_f_ is the inactivation time constant, it also determines the recovery of the channel in the LR1 model) is a major parameter determining T_EAD_.

To more systematically explore the dependence of T_EAD_ on ion channel conductance, we carried out simulations by randomly drawn the maximum conductance of different ionic currents and measured all T_EAD_. The data is presented as a histogram in Fig.6F. The distribution shows that T_EAD_ is mainly between 250 ms and 500 ms (peak ~280 ms), which is the same range seen in Figs.6 A-D (the range is narrower comparing to Fig.6D since the control τ_f_ was used for all the random parameter sets).

Therefore, based on the theoretical predictions and simulation results of the LR1 model, we conclude that the basic T_EAD_ is mainly determined by the time constant of I_Ca,L_ (I_si_ in the LR1 model), which is the oscillation period at the Hopf bifurcation. The change in an ionic current conductance exhibits small effects on T_EAD_ until it causes an EAD to disappear or appear at which T_EAD_ is mainly influenced by the homoclinic bifurcation.

#### Inter-EAD interval in physiologically more detailed models

The simulation results from the physiologically more detailed models show similar characteristics of T_EAD_ dependence on different parameters (Fig.7). Increasing an inward current decreases T_EAD_ while increasing an outward current (except I_to_) increases T_EAD_. In the LRd and H_UCLA_ models, increasing I_to_ conductance decreases T_EAD_. This is because Ito promotes EADs in these models. Similar to the LR1 model, the change in T_EAD_ is not very sensitive to a change in the maximum conductance of an ionic current until it is close to the transition from two EADs to one EAD (or from one EAD to two EADs) in the AP at which T_EAD_ changes rapidly. The predominant T_EAD_ for the 5 models are roughly: LRd—90 ms; H_UCLA_—85 ms; TP04—270 ms; ORd—275 ms; and GB—125 ms. As shown in Fig.7, T_EAD_ in the LRd, H_UCLA_, and GB models are shorter than 200 ms (between 50 ms to 200 ms), while those in the TP04 and ORd models range from 250 ms to 500 ms. In all the models, τf exhibits a stronger effect on T_EAD_ than the other parameters we explored.

**Figure 7.**
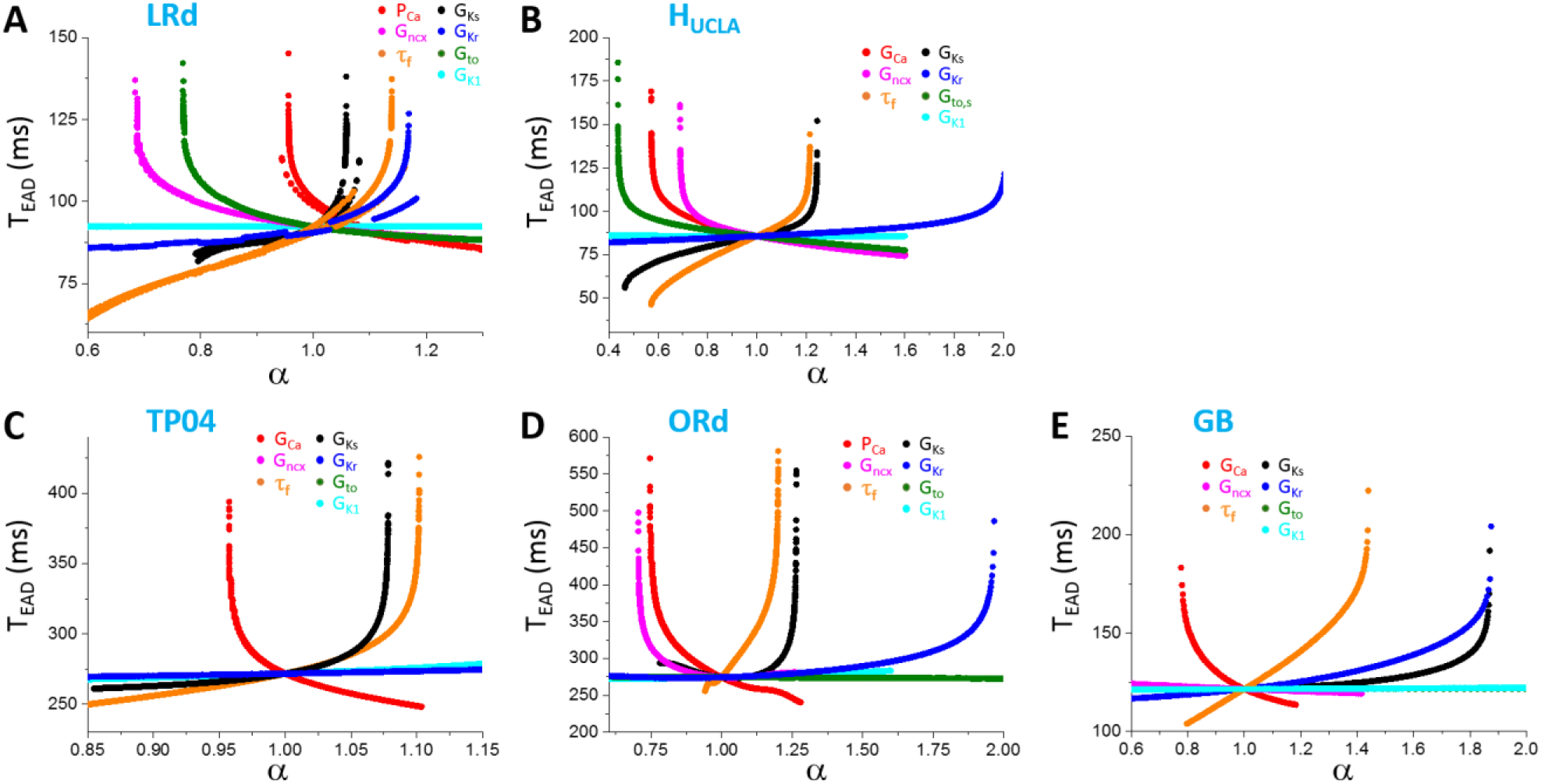
Dependence of T_EAD_ on maximum conductance and kinetics of ionic currents in different models. **A**. LRd. **B**. H_UCLA_. **C**. TP04. **D**. ORd. **E**. GB Only the first T_EAD_ (the interval between the first and the second EAD) are plotted. One parameter (indicated by color) was changed while the others were set at their control values.

In Fig.7, we only show the first inter-EAD interval versus a specific parameter. To explore wider ranges of T_EAD_ in these models, we used random parameter sets (see Methods) and measured all T_EAD_ in an AP. Fig.8 shows the T_EAD_ distributions for the AP models. The T_EAD_ ranges are: LRd—from 75 ms to 125 ms (peak ~90 ms); H_UCLA_—from 50 ms to 150 ms (peak ~85 ms); TP04—from 225 ms to 350 ms (peak ~250 ms); ORd—from 200ms to 500 ms (peak ~250 ms); and GB—from 100 ms to 200 ms (peak ~125 ms). Therefore, the T_EAD_ of the TP04 and ORd model is in the experimentally observed range, while those of the GB, LRD, and H_UCLA_ models are too short comparing to the experimental recordings. Note that the T_EAD_ range using the random parameter sets is similar to the range seen in Fig.7 for each model, indicating that the T_EAD_ range of an AP model is not sensitive to ionic current conductance but mainly determined by the period variation in the dual Hopf-homoclinic bifurcation.

**Figure 8.**
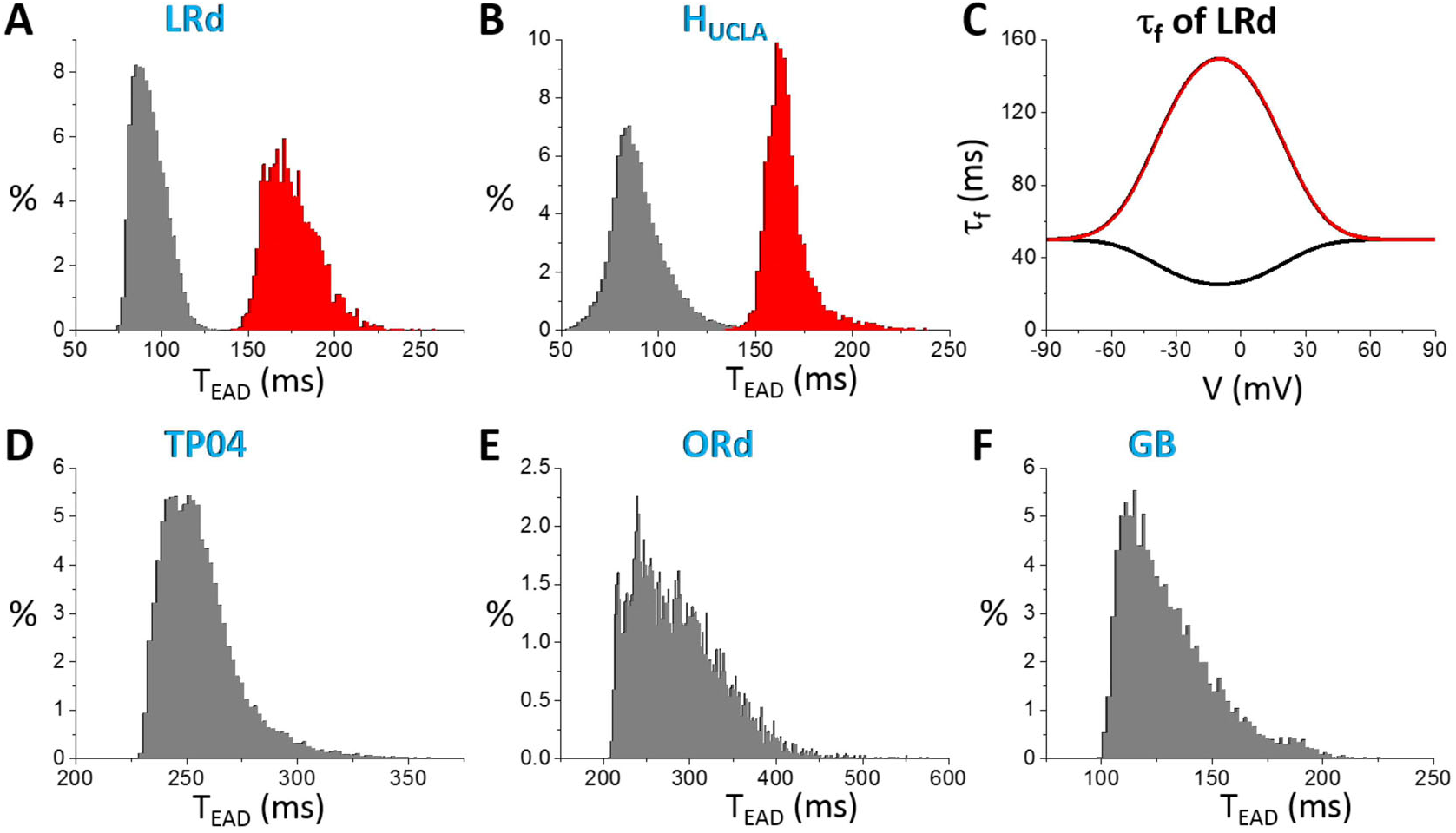
Distributions of T_EAD_ in different models. **A**. LRd. **B**. H_UCLA_. **D**. TP04. **E**. ORd. **F**. GB. In these distributions, the ionic conductance (see SI for the specific ones for each model) were randomly drawn from the assigned intervals. **C**. τ_f_ versus V in the original LRd (black) and the modified one (red). The same τ_f_ was used in the H_UCLA_ model as in the LRd model.

Based on the theoretical predictions that τ_f_ is the main determinant of the inter-EAD interval, we changed τ_f_ functions in the LRd and H_UCLA_ models from upward bell-shaped functions to downward bell-shaped functions (see Fig.8C) to lengthen τ_f_ in the plateau phase, we can effectively shift the T_EAD_ distributions toward the longer periods (red histograms in Figs.8 A and B). The inactivation time constants of I_Ca,L_ in the plateau voltage for the TP04 model and the ORd model are much longer, and thus inter-EAD intervals of these two models are also much longer. The GB model has a much shorter inactivation time constant of I_Ca,L_ in the plateau, similar to those of LRd and H_UCLA_, and thus the inter-EAD interval is also short.

### Determinants of EAD latency

As shown in Table I, L_EAD_ varied in a large range, from less than 100 ms to a couple of seconds. Based on the bifurcation theory of EADs [4,10], for the EADs to occur, besides the instability leading to oscillations, the voltage needs to decay into the window voltage range of I_Ca,L_ activation and the LCCs need to be recovered by a certain amount so that there are enough LCCs available for reopening.

To reveal how L_EAD_ is determined by the ion channel properties, we started with simulations of the LR1 model (Fig.9A). We varied four parameters: G_si_, G_K_, G_K1_, and τ_f_. Increasing G_si_ first decreases L_EAD_ quickly and then increases L_EAD_ slowly (red curve in Fig.9A). The longest L_EAD_ occurs when the first EAD appears in the AP (green arrow in Fig.9A). When a new EAD first appears in an AP, its takeoff potential is the lowest which is close to the potential of the homoclinic bifurcation point (see Fig.2A), and it takes a longer time for the EAD depolarization to occur (the same reason that the T_EAD_ is the longest and A_EAD_ is the largest when a new EAD appears). As G_si_ increases, the takeoff potential is higher and thus L_EAD_ shortens. However, increasing G_si_ also slows the decay of voltage into the window range of LCC reactivation, and thus lengthens L_EAD_. Increasing G_K_ does the opposite (blue curve in Fig.9A) for the same reasons mentioned for G_si_. Changing G_K1_ has no effect since I_K1_ is almost negligible in early phase-2 of the AP. τ_f_ has a big effect on L_EAD_, which can vary L_EAD_ in a much wider range than the conductance. τ_f_ affects L_EAD_ by two ways: 1) slowing τ_f_ causes a slower inactivation of Isi which delays the voltage decay to the window range; 2) slowing τ_f_ delays recovery of LCCs, which then delays the depolarization of the first EAD.

**Figure 9.**
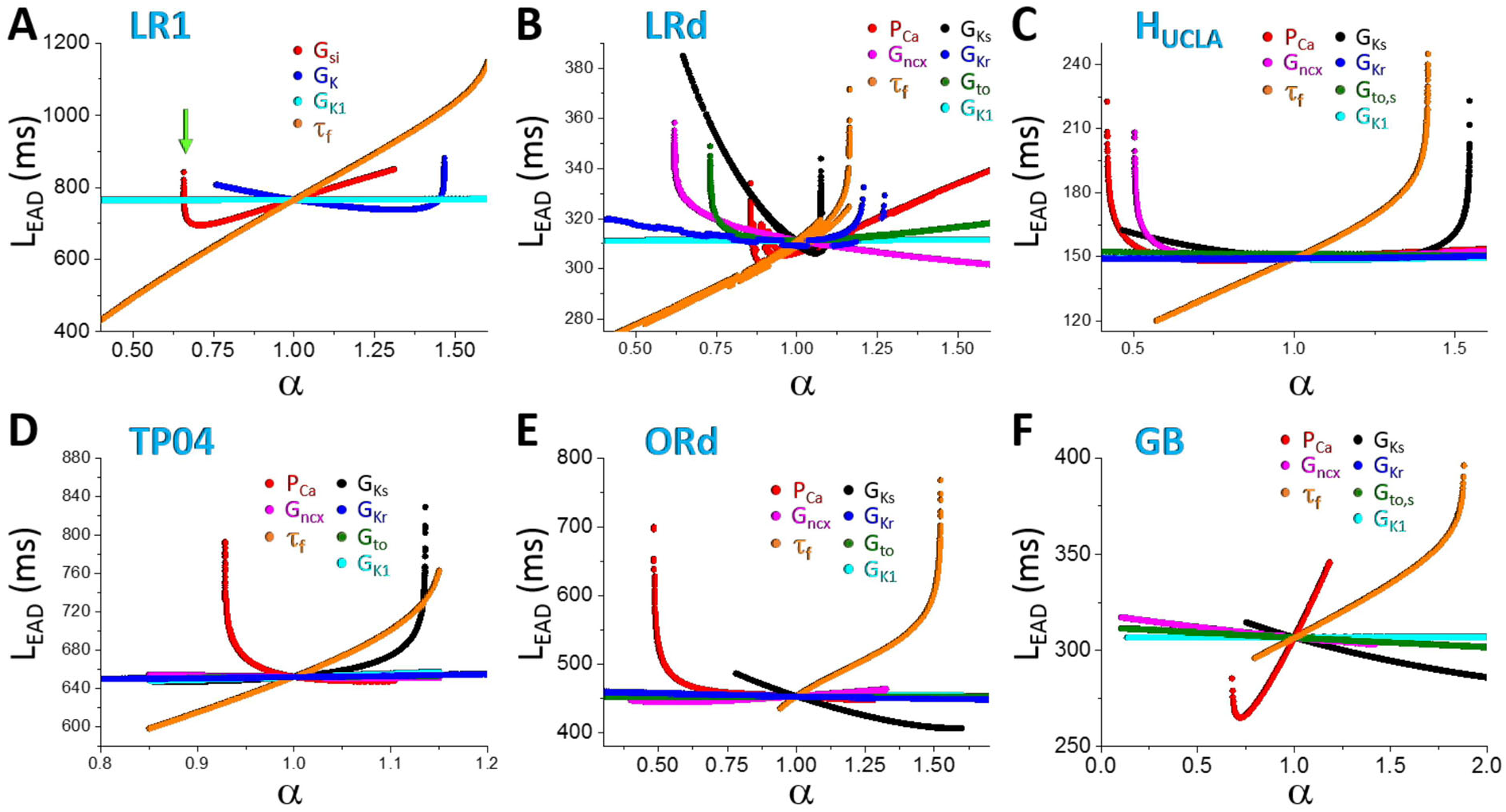
Dependence of EAD latency on ionic current conductance in different AP models. **A**. LR1. **B**. LRd. **C**. H_UCLA_. **D**. TP04. **E**. ORd. **F**. GB. One parameter (indicated by color) was changed while the others were set as their control values.

Similar behaviors occur in all other models (Figs.9B-F): changing a conductance usually has a small effect on L_EAD_ until it causes the only EAD in the AP to disappear, at which L_EAD_ changes steeply; and the inactivation time constant of I_Ca,L_ is the most sensitive parameter for L_EAD_. The L_EAD_ varies largely from model to model. In Fig. 10, we show L_EAD_ distributions from random parameter sets for all models, and the L_EAD_ ranges are: LR1—from 600 ms to 1000 ms (peak ~750 ms); LRd—from 240 ms to 550 ms (peak ~300 ms); H_UCLA_—from 135 ms to 180 ms (peak ~150 ms); TP04—from 640 ms to 720 ms (peak~650 ms); ORd—from 360 ms to 600 ms (peak ~400 ms); GB—from 240 ms to 500 ms (peak~280 ms).

**Figure 10.**
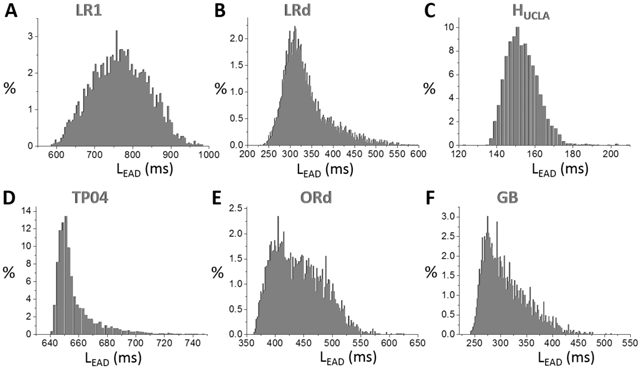
Distributions of T_EAD_ in different models. **A**. LR1. **B**. LRd. **C**. H_UCLA_. **D**. TP04. **E**. ORd. **F**. GB. In these distributions, the ionic conductance (see SI for the specific ones for each model) were randomly drawn from the assigned intervals.

## DISCUSSION

In this study, we used theoretical analyses and computer simulations to investigate EAD properties and their major determinants in AP models of ventricular myocytes. Our major observations and conclusions are summarized and discussed below.

### EAD takeoff potential and amplitude

In all the models we simulated, the EAD takeoff potentials are usually above −40 mV (see the 0 mV shift cases in Fig.3), which is in agreement with experimental data from isolated ventricular myocytes (Table I). A negative linear correlation between EAD amplitude and takeoff potential, which has been shown in experimental recordings [1-3], can be implied from the dual Hopf-homoclinic bifurcation and is shown in simulations of all models. The slopes of the negative linear correlations, the lowest takeoff potentials, and the maximum EAD amplitudes vary substantially from model to model. The ORd model exhibits the largest maximum EAD amplitude and the steepest slope of the linear correlation between EAD amplitude and takeoff potential.

Although EADs are promoted by increasing inward currents or decreasing outward currents, once EADs occur in an AP, increasing the maximum conductance of an inward current or decreasing that of an outward current does not increase the amplitudes of the EADs. Increasing the maximum conductance of an inward current causes more EADs in the AP, and the maximum EAD amplitude occurs when a new EAD appears in the AP. The amplitude of this new EAD then decreases as the conductance increases. Increasing the maximum conductance of an outward current decreases the number of EADs in the AP but increases the EAD amplitude. The maximum EAD amplitude occurs before an EAD disappears from the AP. Based on the bifurcation analysis [4, 14-16], the EAD amplitude grows as the system evolves from the Hopf bifurcation point to the homoclinic bifurcation point (Fig.2A and Fig.4). This behavior should hold for both supercritical [4,16] and subcritical [15,16] Hopf bifurcation. At the homoclinic bifurcation point, the takeoff potential is the lowest and the EAD amplitude becomes the maximum. Therefore, the amplitude of the last EAD in an AP depends on how far away the takeoff potential is from the homoclinic bifurcation point. Born of a new EAD or death of an existing EAD always occurs when the EAD takes off near the homoclinic bifurcation point.

Note that in our simulations of assessing EAD amplitude and takeoff potential, we used random parameter sets to explore a wide range of parameters for each model. In these simulations, only phase-2 EADs in the ventricular myocyte models were observed. A recent simulation study [16] using the ORd model and the Kurata et al model [54] also showed that only phase-2 EADs could be observed. This indicates that varying maximum ionic current conductance may not produce phase-3 EADs. This agrees with the experimental data that EADs observed in isolated ventricular myocytes are mainly phase-2 EADs (Table I), while phase-3 EADs are rarely observed in isolated cells, except under Ca^2+^ overload [55] or by external current injection [47]. On the other hand, phase-3 EADs are observed in Purkinje fibers [17] and cardiac tissue [56-58]. Previous computer simulations showed that phase-3 EADs could occur in single cells with a strong I_nsCa_ under elevated intracellular Ca^2+^ concentration [18,19,59] or in tissue with repolarization heterogeneity induced dynamical instabilities [20,56]. Therefore, phase-3 EADs can be caused either by strong Ca^2+^ overload in isolated myocytes or by repolarization heterogeneities in tissue with reduced repolarization reserve, while phase-2 EADs are mainly due to reduced repolarization reserve and reactivation of I_Ca,L_ [2,10,12,60,61].

Since phase-2 EADs cannot propagate into PVCs in tissue [17-20], this raises question on how are EADs linked to arrhythmias under LQTS and many other diseased conditions where Ca^2+^ may not be overloaded. In recent studies [20,62], we demonstrated how phase-2 EADs and tissue-scale dynamical instabilities interact to result in PVCs and arrhythmias under LQTS, linking mechanistically phase-2 EADs to arrhythmogenesis.

### Inter-EAD interval

Based on the bifurcation analysis, the inter-EAD interval is governed by the period of the limit cycle oscillation between the Hopf bifurcation and the homoclinic bifurcation. Similar to A_EAD_, the inter-EAD interval increases as the system evolves from the Hopf bifurcation point to the homoclinic bifurcation point. This behavior has been demonstrated in experimental recordings previously [63]. Our theoretical analysis and simulation of the AP models showed that the inter-EAD intervals (except the last one in an AP), which are mainly determined by the period of the limit cycle oscillation at the Hopf bifurcation, is insensitive to the change of maximum ionic current conductance but more sensitive to the recovery of LCCs (Figs.7 and 8, and Eq.7). The T_EAD_ range of an AP model is determined by the oscillation period between the Hopf bifurcation and the homoclinic bifurcation. However, the inter-EAD interval from different models exhibits different ranges, differing several folds. On the other hand, the inter-EAD intervals observed in isolated ventricular myocyte experiments mostly are in the range from 200 ms to 500 ms, irrespective of species (Table I). In the AP models simulated in this study, the inter-EAD intervals of LR1, TP04, and ORd are in the same range as observed experimentally, but other models exhibited much faster inter-EAD intervals, indicating that caveats may exist in these models. Our simulation indicates that the major caveat may lie in the formulation of I_Ca,L_, in particular the recovery time of I_Ca,L_ during the plateau phase (see more detailed discussion below).

### EAD latency

EAD latency is determined by many factors. In term of biophysics, since EADs are caused by reactivation of LCCs, the voltage needs to decay into the window range for reactivation of LCCs, which depends on the speed of activation of outward currents (namely, I_Ks_, I_Kr_, and I_to_) and inactivation of inward currents (namely I_Ca,L_). Then there are enough LCCs recovered for re-opening, which depends on how fast the LCCs recovers. In more general term of nonlinear dynamics as indicated by the bifurcation theory, the voltage and other variables need to enter the basin or the vicinity of the basin of attraction of the limit cycle. This requires not only the voltage but also the other state variables to reach their proper values. For example, in the LR1 model, the X-gating variable needs to grow to a certain value to engage the Hopf bifurcation as shown in Fig.2A. If X grows too slowly, the system may stay at the quasi-equilibrium state for a long time with no oscillations until reaching the Hopf bifurcation point, such as the cases shown in Song et al [49]. Note that transient oscillations around a stable focus can occur before the Hopf bifurcation, and thus EADs can occur before the Hopf bifurcation (see Fig.2A, Figs.4 C and D in this study, and Figs.2 and 4 in the study by Kügler [15]). Therefore, the EAD latency can be very variable, explaining the experimental observation that EAD latency varies in a wide range, from less than 100 ms to several seconds (Table I).

### Implications to mathematical modeling of cardiac APs

It is obvious that understanding the EAD properties and nonlinear dynamics is of great importance for understanding arrhythmogenesis in cardiac diseases [64], but it also provides important information for cardiac AP modeling. Previous AP modeling has mainly considered AP morphology, APD, as well as APD restitution, but not the EAD properties. For example, the inter-EAD intervals of the guinea pig ventricular myocyte models [13,21,22] are much shorter than what have been observed in isolated guinea pig ventricular myocytes (Table I). This is also true for the rabbit ventricular myocyte models. As shown in Figs. 8 A and B, we can effectively increase the inter-EAD interval by increasing the inactivation time constant τ_f_ of I_Ca,L_. However, for both the guinea pig model [65] and the rabbit model [66], the original inactivation time constants were based on experimental measurements. Since in the Hodgkin-Huxley formulation of I_Ca,L_, the f-gate is a voltage-dependent inactivation gate, but it also governs the recovery of I_Ca,L_. Therefore, one would conclude that experimentally-based τ_f_ might be a correct constant but the recovery properties of I_Ca,L_ in these models are incorrect, which gives rise to the discrepancy in inter-EAD intervals between mathematical models and experimental measurements. On the other hand, τ_f_ is large in the LR1, TP04, and ORd models, which give rise to the right recovery times to result in inter-EAD intervals in the ranges as observed in experiments. That also does not mean that I_Ca,L_ models are completely correct in these AP models since we know that the one for the LR1 model gives rise to a too slow inactivation of I_Ca,L_. Therefore, the EAD properties provide additional important information for AP modeling, which need to be considered in future AP model development.

### Limitations

A major limitation is reliable experimental data curation. First, most of the values in Table I were estimated from the published figures, which is difficult to be accurate and unbiased. Second, experimental data of EADs recorded from isolated ventricular is not abundant. Moreover, to calculate T_EAD_, we have to select APs with two or more EADs, which further limited our data sources. Third, most of the experimental plots do not have coordinates but indicated by scale bars. Sometimes, these scale bars may not be correctly labeled due to different reasons. For example, we confirmed with Dr. Li that the time scale bar in Fig.2C of Ref. [31] is 300 ms instead of 150 ms. In computer simulations, we only explored the maximum conductance of the ion currents and the inactivation time constant of I_Ca,L_, but it is obvious any parameter that affects repolarization will have an effect on EAD properties. For example, the time constants of ion channel activation and inactivation, the Ca^2+^-dependent inaction of I_Ca,L_, the intracellular Ca^2+^ transient, as well as spatial distribution of the ion channels will impact the EAD behaviors, which need to be investigated in future studies. Another limitation of the simulations is that our conclusions may depend on the setting of control parameter and the assigned intervals for random parameters. However, as we show in this study, despite certain distinct difference between models, such as the inter-EAD interval, the general conclusions are not model dependent, and thus, not likely to be affected by the choice of control parameter sets and the assigned intervals.

## AUTHOR CONTRIBUTIONS

Z.Q. conveyed the overall design and wrote the manuscript; X.H. designed and performed the computer simulations; Z.Q. and Z.S. curated experimental data. All authors participated in interpreting the results and edited the manuscript.

